# Integrating theory and machine learning to reveal determinants of plasmid copy number

**DOI:** 10.1101/2025.10.23.684078

**Authors:** Iqra Shahzadi, Wenzhi Xue, Hasan Ubaid Ullah, Rohan Maddamsetti, Lingchong You, Teng Wang

**Affiliations:** State Key Laboratory of Quantitative Synthetic Biology, Shenzhen Institute of Synthetic Biology, Shenzhen Institutes of Advanced Technology, Chinese Academy of Sciences, Shenzhen, China; University of Chinese Academy of Sciences, Beijing, China; Department of Biochemistry and Microbiology, Rutgers University, New Brunswick, NJ, USA; Center for Quantitative Biodesign, Duke University, Durham, NC, USA; Department of Biomedical Engineering, Duke University, Durham, NC, USA; Department of Molecular Genetics and Microbiology, Duke University School of Medicine, Durham, NC, USA

## Abstract

Plasmids are extrachromosomal mobile genetic elements whose copy numbers (PCNs) critically influence microbial evolution, antibiotic resistance and pathogenicity. Despite their importance and immense diversity, the ecological, evolutionary and molecular factors determining PCN remain poorly understood. Here, we present a theoretical model to explain the empirical power-law relationship between plasmid size and copy number, one of the fundamental quantitative principles governing PCN control. However, this relationship alone has limited predictive power. To improve PCN prediction, we introduce a data-driven approach incorporating diverse features. Trained on >10,000 plasmids, our machine learning model achieves significantly enhanced accuracy, with plasmid-encoded protein domains emerging as key predictors. Applying this framework, we conduct the first comprehensive analysis of PCN distributions across hundreds of thousands of metagenomic plasmids (IMG/PR database) and tens of thousands of clinical isolates, uncovering niche specific taxonomic PCN hotspots and ecological adaptations. These results provide critical insights into plasmid ecology, ARG surveillance and shed lights on the gut plasmidome, a “dark matter” in human microbiome.

## Introduction

Plasmids are extrachromosomal mobile genetic elements that play critical roles in microbial genomics, ecology and evolution^1^. Plasmids typically exist in multiple copies per cell, with copy numbers (PCNs) spanning several orders of magnitude^2,3^. PCN holds significance in multiple scenarios, from environment, health to engineering. Multicopy plasmids can potentiate bacterial evolution, promoting the emergence and dissemination of antibiotic resistance^4–6^. Many virulence factors are plasmid-encoded, with PCN variation playing a pivotal role in the pathogenesis of microbes such as *Yersinia pestis*, the plague bacterium^7–9^. In genetic engineering, plasmids serve as invaluable tools, where PCN directly determines the gene dosage, expression level, metabolic burden and circuit stability^10–13^.

Natural environments harbor a vast diversity of plasmids, evidenced by repositories like the global IMG/PR database which currently houses nearly 700,000 plasmid sequences from different ecosystems from the human gut to the ocean^14^. By contrast, experimental measurements of PCN (e.g., via qPCR or Southern blot) remain labor-intensive and low-throughput^15^. While high-throughput sequencing enables PCN estimation through coverage ratio analysis (plasmid versus chromosomal reads), current pipelines have only characterized PCNs for about 12,000 plasmids - a small fraction of natural plasmid diversity - leaving the copy numbers of most plasmids unknown^2,3^.

This striking disparity highlights the necessity of computational PCN predictions. Large-scale PCN predictions will help deduce the quantitative principles and ecological patterns governing copy number variations. Such predictive capability is particularly valuable in environments like human gut, where the uncultivability of many microbial organisms makes direct PCN measurement experimentally challenging^16^. Moreover, given the growing antibiotic resistance crisis, reliable PCN prediction would enable precise assessment of plasmid-borne resistance gene dosage, thereby improving our ability to evaluate and control dissemination risk^17^. Additionally, accurate PCN prediction would support the rational design of synthetic gene circuits, facilitating the development of predictable synthetic biology systems^18^.

Previous studies identified an inverse power-law relationship between plasmid length and copy number, one of the fundamental rules governing PCN^2,3^. Here, we first developed a theoretical model to explain this empirical relationship, confirming plasmid size as an important PCN predictor. However, this relationship alone has limited predictive power. To overcome this limitation, we designed a machine learning framework incorporating diverse plasmid features, which significantly improved prediction accuracy and highlighted the role of plasmid-encoded protein domains in shaping PCN. Applying this model to clinical plasmids revealed the quantitative interplay between PCN and ARG dosage. We further extended this framework to the IMG/PR dataset, uncovering ecosystem specific PCN trends, particularly in the human gut microbiome. Combining theoretical modeling and data-driven approaches, this work elucidates quantitative principles of PCN control and offers a framework for precise PCN prediction, which have broad implications for microbial ecology, evolution, environment and human health.

## Results

### A simple theory explains how PCN correlates with plasmid size

To uncover the quantitative principles governing PCN regulation, we first analyzed a comprehensive PCN dataset developed in a previous study^3,19^ (Methods). This dataset, encompassing 11,338 prokaryotic plasmids, was built upon 4,317 plasmid-carrying genomes whose short-read sequencing data were provided in the NCBI SRA^20^. For each plasmid, PCN was estimated by calculating its mean sequencing coverage relative to that of its host chromosome, assuming that the abundances of reads mapping to the chromosome or plasmids reflected their physical DNA amounts in the host cell. Note that these PCN estimates represent relative copy numbers (plasmid-to-chromosome ratios), not necessarily absolute plasmid counts per cell^3^. PCN values may fall below one if the cell contains multiple chromosome copies or if plasmids are absent in some cells of the sequenced population^21–24^.

The majority (99.77%) of plasmids in the dataset are from bacteria, while the rest are from archaea. *Escherichia* accounts for the largest proportion (∼20%) of plasmids, followed by *Klebsiella* (∼18%) and Salmonella (∼5%). Plasmid length varied significantly, ranging from 1 kb to 2.59 Mb, with a mean of 94 kb and a median of 55.8 kb. While most plasmids in the dataset come from natural environments, 0.2% are engineered (constructed or modified in laboratories, see Methods). Ecologically, the dataset covers diverse microbial ecosystems, with the human-associated environment contributing the largest proportion of plasmids (37.7%), followed by livestock (7.37%).

This dataset revealed an inverse power law correlation between plasmid length and PCN (Figure 1A), characterized by a scaling exponent of -0.71. This suggests a tradeoff between plasmid length and copy number, where larger plasmids generally maintain lower copy number, a pattern also reported in previous studies^2,3^. Yet the quantitative basis of this power-law relationship remains unclear.

**Figure 1.**
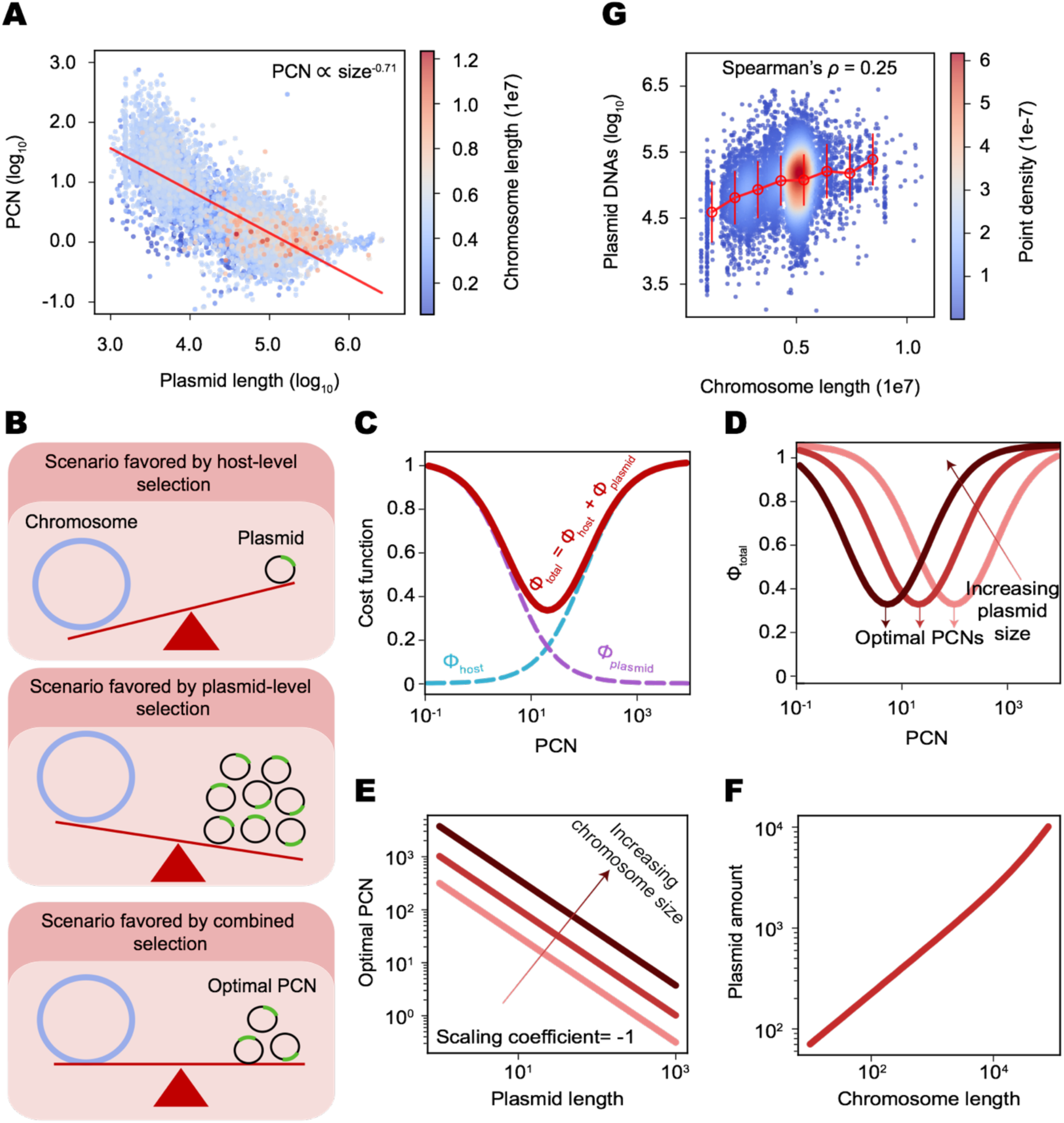
Multi-level selections explain the power-law relationship between plasmid size and PCN. (A) The inverse power-law relationship between plasmid size and PCN, with chromosome length indicated by color gradient. Larger plasmids generally have lower copy numbers, reflecting an evolutionary size-PCN tradeoff. (B) The plasmid size-PCN tradeoff arises from the interplay between two selective forces. Host-level selection favors plasmids with minimal metabolic burden (top panel), while plasmid-level selection favors those with maximized self-replication and transfer (middle panel). The optimal PCN emerges from the interplay of these two opposing forces (bottom panel). (C) Theoretical curves showing the cost functions for host-level selection (𝜙*_host_*, blue dashed line) and plasmid-level selection (𝜙*_plasmid_*, purple dashed line). Selection at each level operates by minimizing its respective cost function. Because these objectives are inherently conflicting, the optimal PCN emerges as the value that minimizes the combined cost (𝜙*_total_* = 𝜙*_host_* + 𝜙*_plasmid_*, red solid line). 𝜙*_host_* was formulated as 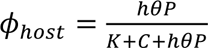, where *C*, *P* and 𝜃 represent chromosome size, plasmid size and PCN, respectively. 𝐾 is a constant characterizing how the total amount of intracellular resources changes with genome size. *K* is a constant describing the relative advantage of plasmid genes in resource competition. 𝜙*_plasmid_* was formulated as 𝜙*_plasmid_* = 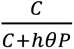. Here the theoretical curves were generated with *K* = 5 × 10^4^, 𝐶 = 2000, *P* = 50 and *h* = 10. Varying parameters did not change the shapes of the curves. (D) The relationships between 𝜙*_total_* and PCN, under different plasmid sizes. There different plasmid sizes (𝑃 = 10, 50 and 200 from right to left) were tested. Other parameters are 𝐾 = 5 × 10^4^, 𝐶 = 2000 and ℎ = 10. (E) Theoretically predicted plasmid size-PCN tradeoffs under three different chromosome sizes. Here, the scaling exponent between plasmid size and PCN equals -1. Three different chromosome sizes (𝐶 = 2 × 10^2^, 2 × 10^3^ and 2 × 10^4^ from left to right) were tested with 𝐾 = 5 × 10^4^ and ℎ = 10. (F) Theoretically predicted positive correlation between chromosome length and plasmid DNA amount. Here plasmid DNA amount was calculated as plasmid size multiplied by its PCN. Other parameters are 𝐾 = 5 × 10^4^ and ℎ = 10. (G) Observed positive correlation (Spearman’s *ρ* = 0.25) between chromosome length and plasmid DNA amount, with point density represented by color shading.

From first principles, the persistence of a plasmid in microbial genomes was shaped by two levels of selective forces: host-level selection and plasmid-level selection within a host (Figure 1B)^25,26^. Host-level selection tends to eliminate plasmids that impose high metabolic burden. This occurs because plasmid gene expression consumes energy and resources in host cell, reducing host fitness and causing plasmid-bearing cells to be competitively excluded by plasmid-free ones. However, even burdensome plasmids may persist by exploiting strategies such as highly efficient replication or horizontal transfer^27,28^. These plasmid-centric forces, collectively referred to as plasmid-level selection^25,26^, encompass all mechanisms that increase plasmid persistence at host’s expense, including plasmid replication rate, transfer efficiency, and reduced segregation error. Long-term plasmid stability in prokaryotic genomes thus emerges from the interplay of these opposing forces.

To explain the quantitative tradeoff between plasmid size and PCN, we developed a mathematical model in which host- and plasmid-level forces are formalized as outcomes of intracellular resource competition between chromosomes and plasmids^25^ (see Supplementary Information for details). In this framework, plasmid competitiveness scales with its total DNA content (plasmid size × PCN). We then defined the cost functions for host-level (𝜙*_host_*) and plasmid-level (𝜙*_plasmid_*) forces, which could be derived analytically as Hill functions of chromosome size and plasmid DNA content (Figure 1C). Selection at each level operates by minimizing its respective cost function. Because these objectives are inherently conflicting, the optimal PCN emerges as the value that minimizes the combined cost function (𝜙*_total_*), defined as the sum of 𝜙*_host_* and 𝜙*_plasmid_*(Figure 1C). In this way, plasmid evolution is treated as an optimization problem. Owing to the model’s simplicity, the optimal PCN can be solved analytically as a function of both chromosome and plasmid size (Figure 1D).

The theory successfully predicts the power-law relationship between plasmid size and PCN, albeit with a scaling exponent of -1 (Figure 1E). The discrepancy between the predicted and observed scaling exponents may be due to the model’s simplicity. For instance, the model didn’t account for the functional heterogeneity of chromosome or plasmid-encoded genes. Additionally, it overlooked specific gene conflicts between plasmids and chromosomes, an important source of plasmid fitness burdens^29^. Nevertheless, the model still predicts a positive correlation between plasmid DNA content and chromosome size (Figure 1F), which was confirmed by PCN dataset analysis (Spearman’s 𝜌 = 0.25; Figure 1G, Supplementary Figure S1A)^2^. These findings provide a mechanistic basis for understanding the copy number control of prokaryotic plasmids.

While our theory provides a plausible explanation for the tradeoff between PCN and plasmid size, its quantitative predictive power remains limited (Figure 1G). The power law fit is also inadequate (*R*² ∼0.63, Supplementary Figure S1B), as it ignores other factors influencing copy number regulation, especially plasmid-encoded functions. This gap underscores the necessity for a data-driven approach to model PCN that considers genetic features beyond plasmid length.

### PCN prediction through a multi-feature machine learning framework

To improve prediction accuracy, we developed a random forest regression model trained on diverse genomic and plasmid-level features, with a strategic emphasis on plasmid-encoded protein domains. This domain-centric approach addresses a key limitation in plasmid biology: many plasmid-encoded proteins lack functional annotations in public databases and are simply labeled as ‘hypothetical’^30^. Traditional analyses relying on annotated proteins risk overlooking these uncharacterized elements^31^, biasing predictions toward well-studied systems and limiting their applicability in many real-world scenarios like metagenomes, where functional annotations can be sparse. By contrast, protein domains, detectable even in unannotated sequences, represent evolutionary conserved functional units^32^. This framework ensures that our model remains robust across diverse datasets, including poorly annotated plasmids from clinical or environmental isolates^14^.

We first used Prodigal to identify protein-coding sequences (CDSs) of all plasmids^33^. Domain annotation of these CDSs against the Pfam database yielded 11,533 unique Pfam domains^32^ (Methods). To find domains relevant to PCN, we first performed a point-biserial correlation analysis across all domains, followed by multiple testing correction (Benjamini– Hochberg FDR, q < 0.05). This analysis revealed 1,288 (∼9%) domains significantly associated with PCN, indicating their potential roles in copy number regulation (Figure 2A). We created binary features for each plasmid based on the presence or absence of these domains. Besides domains, we also incorporated other features in our model, including plasmid length, host chromosomal length, and plasmid *k*-mer frequencies (up to 3-mers), to construct a multi-modal feature matrix that captures both sequence composition and functional properties.

**Figure 2.**
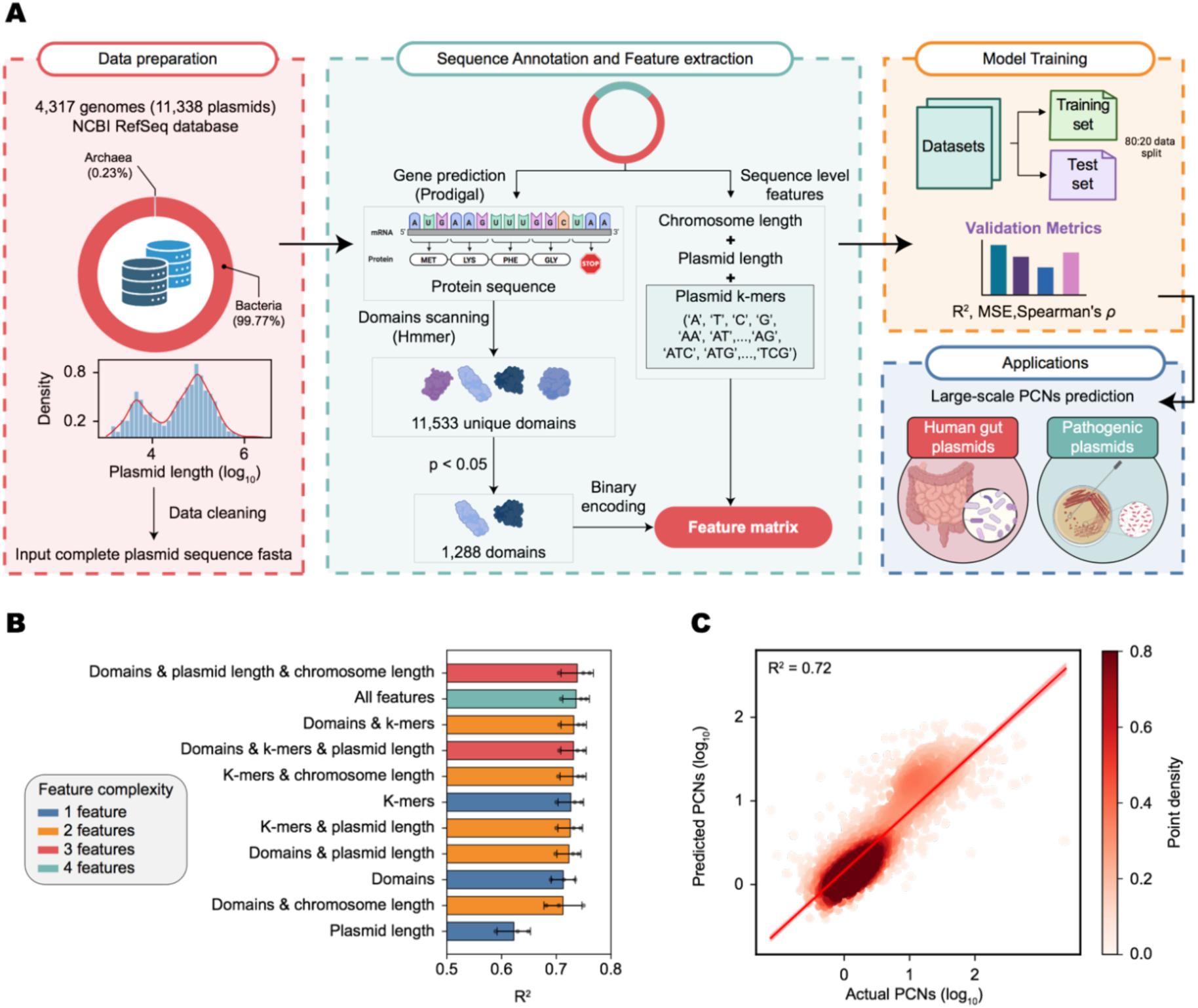
A machine learning framework that predicts plasmid copy number (PCN) using sequence-derived features. (A) The computational pipeline. We retrieved 11,338 plasmid sequences from 4,317 genomes (99.77% bacterial, 0.23% archaeal). Protein domains were annotated using HMMER (Pfam), with 1,288 domains showing significant correlation with PCN. Features (plasmid length, chromosome length, *k*-mers, plasmid encoded domains) were combined into a matrix. The dataset was split at a 4:1 ratio for training and test, and performance was evaluated with R², MSE, and Spearman’s *ρ*. (B) Multi-feature models (domains, *k*-mers, replicon lengths) achieved higher R² values, outperforming single-feature models. Error bars represent the standard deviations across 3 replicates. (C) The correlation between actual PCN values and those predicted by the model using all features. Point density is represented by red shading (darker indicates higher density).

To assess the impact of feature selection on predictive accuracy, we systematically tested various feature combinations. For each combination, the model was trained and evaluated through 3 independent training-test splits at 4:1 ratio (Methods). The results showed that models integrating multiple feature types consistently outperformed single-feature models, with the highest accuracy achieved when combining domains, plasmid length, and chromosomal length (Figure 2B; Supplementary Figure S2). Consistent with our mechanistic modeling of the power-law scaling, plasmid length alone had only moderate predictive power, reinforcing the necessity of a multi-feature approach for reliable PCN estimation (Figure 2B; Supplementary Figure S2).

Given the different availability of plasmid host information in various applications, we chose two models for different use cases. The first model (*R*² ∼0.72) incorporates plasmid domains, *k*-mers, plasmid length and chromosomal length, making it suitable for studies with accessible host genome data (Figure 2C). However, in scenarios like metagenomics, host chromosomal information is often unavailable. To address this, we chose a simplified, chromosome-independent model using only plasmid-derived features (domains, plasmid *k*-mers, and plasmid length), which maintained high accuracy (*R*² ∼0.71). This flexibility ensures broad applicability, enabling PCN prediction even in datasets where plasmid hosts are unknown. Together, these models provide a powerful framework for PCN prediction in diverse contexts.

### Associations between PCN and plasmid-encoded protein domains

Evolutionary related protein domains can be organized into broader groups known as clans^34^. Domains within a clan typically share sequence, structural or functional similarities. Among the domains significantly associated with PCN, the most prevalent clans are P-loop NTPase, helix - turn - helix (HTH), and NADP_Rossmann, together accounting for 27% of all identified domains (Figure 3A). Notably, these clans are significantly enriched relative to their baseline abundances across all plasmid - encoded domains (Figure 3B). The P-loop NTPase clan, mainly composed of Type III restriction subunits (PF04851) and FtsK/SpoIIIE family proteins (PF01580), drives ATP-dependent DNA processing (GO:0005524)^35^. It performs critical replication and segregation functions through helicase activity and strand translocation (GO:0003677). Meanwhile, HTH domains, especially Sigma-70 factors (PF08281) and HTH AraC-family regulators (PF00165), regulate gene expression adaptively via sequence-specific DNA binding (GO:0043565) and transcription initiation (GO:0006352), allowing plasmids to dynamically respond to antibiotic pressure or host environmental shifts^36–38^. The NADP_Rossmann clan, rich in oxidoreductases like 3-beta hydroxysteroid dehydrogenase (3Beta_HSD) (PF01073), sustains plasmid viability under stress by mediating NADP-dependent redox reactions (GO:0016616) and metabolic integration with host pathways^39^. Together, these clans form an optimized functional triad: P-loop NTPases ensure faithful inheritance, HTH domains adjust gene expression for rapid adaptation, and NADP_Rossmann enzymes maintain metabolic compatibility in various niches. This architecture underscores how plasmids balance replication fidelity, transcriptional flexibility and physiological resilience to thrive as vectors of horizontal gene transfer.

**Figure 3.**
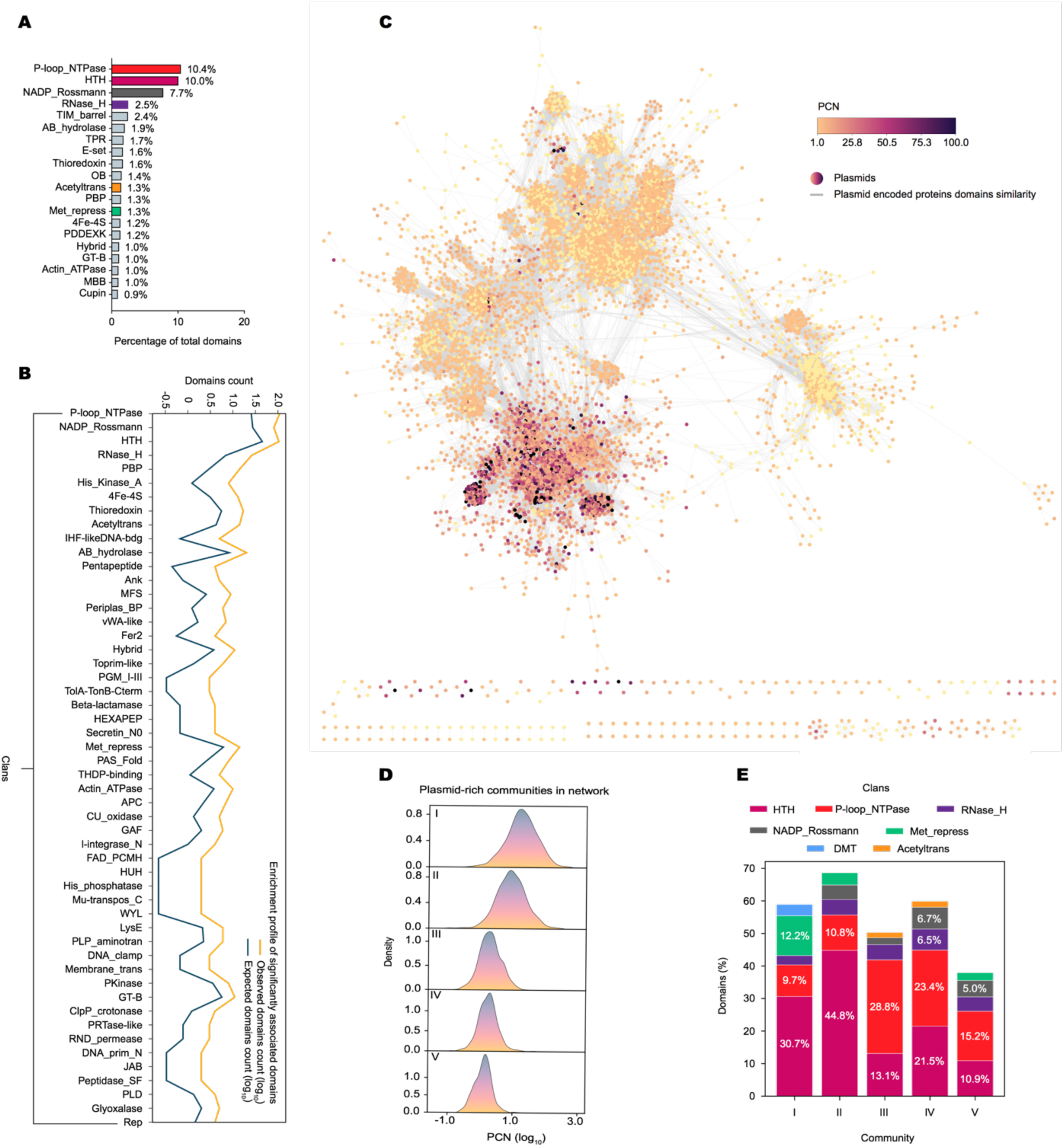
PCN was shaped by plasmid-encoded protein domains. (A) Fractions of top 20 clans in the 1,288 protein domains that are significantly associated with PCNs. (B) Enrichments of different clans relative to their baseline abundances across all plasmid-encoded domains. The expected domain count in each clan was calculated under the null hypothesis of random distribution using the following parameters: the total number of selected domains (1,288), the total number of domains in the dataset (11,533), and the total number of domains in each clan. Statistical significance of observed vs expected distributions was calculated by hypergeometric tests (p< 0.05 threshold). Only clans showing significant enrichment are shown here. (C) Domain similarity network of 10,707 plasmids, colored by PCN (darker indicates higher PCN), with edges representing cosine similarity (cutoff = 0.5) in domain composition. (D) PCN distribution across different plasmid communities. Kernel density estimation plots of PCNs (logarithmic scale) in the five major plasmid clusters are shown. (E) Functional specialization of plasmid communities. Stacked bar plot showing clan composition across communities.

Plasmid-encoded domains allow us to construct a network that visualizes the functional similarities among plasmids. For every plasmid pair, we calculated the cosine similarity of their encoded domains (Methods). After applying a stringent similarity threshold (cosine similarity ≥ 0.5), we created a network with 10,707 nodes (each representing a plasmid) and 1,039,492 edges connecting functionally related plasmids. This network allows us to examine the distribution patterns of plasmid functions and the relationships between plasmid functional content and copy number (Figure 3C, Supplementary Figure S3A).

In the network, plasmids formed tightly interconnected clusters, and this pattern persisted even under more stringent similarity thresholds (cosine similarity ≥ 0.7; Supplementary Figure S3B). Notably, low-copy (<5 copies per cell) and high-copy plasmids (>20 copies per cell) were located in different regions of the network with little overlap. Applying the Louvain community detection algorithm^40^, 112 functionally distinct clusters were identified within the network (Supplementary Figure S3C). Among them, the five largest clusters (I–V), collectively comprising 4,846 plasmids (45% of the total network), exhibited clear differences in both PCN distributions and domain compositions (Figure 3D and E). Clusters I and II, the high-PCN clusters, were highly enriched in HTH domains (30.7% and 44.8%, respectively), while clusters III, IV and V were dominated by low-copy plasmids with greatest enrichment of P-loop NTPase clans (28.8%, 23.4% and 15.2% respectively) (Figure 3E). Together, these analyses create a comprehensive map of PCN regulation where copy number variation results from functional constraints on conserved protein domains.

### Model based prediction identifies high-copy, ARG-rich plasmids in clinical pathogens

Plasmids are key vehicles of horizontal gene transfer, mediating the spread of ARGs and virulence factors among microbes^1^. Plasmids in prokaryotic pathogens substantially contribute to their drug resistance^41^. Predicting the copy numbers of these clinical plasmids is crucial for understanding the dosage and transfer potential of pathogens’ ARGs and is significant for the surveillance and risk assessment of ARG outbreaks. However, due to the lack of predictive framework, the copy number diversity of pathogenic plasmids remains largely unknown.

To address this, we applied our machine-learning approach to the plasmids from NCBI’s Pathogen Detection database, a comprehensive resource of microbial pathogen genomes from food, environment and patient samples^42^. We retrieved the genomes of all fully sequenced prokaryotic pathogens available as of May 2, 2025. These genomes cover 88 pathogen species, with *S. enterica*, *E. coli* and *Shigella* being the most represented. From the 15,855 genome assemblies obtained, we identified 30,254 plasmids, with ∼67.3% of the genomes carrying at least one plasmid.

We applied the full-context model to predict the copy numbers of all pathogenic plasmids by integrating plasmid-encoded domains, *k*-mers, plasmid length and chromosomal length. The predicted PCNs, reaching up to 128, showed substantial diversity and maintained a power-law relationship with plasmid size, characterized by a scaling coefficient of -0.73 (Supplementary Figure S4A). The close agreement with the coefficient derived from the training dataset (Figure 1A) underscores the robustness of our machine learning model. Moreover, we observed a geographic pattern, where high-copy-number plasmids (PCN >10) were overrepresented in samples from Russia, Sweden and India (Figure 4A, Supplementary Figure S4B). Further stratification by anatomical isolation sites showed that pathogenic plasmids from human genital samples had elevated PCN (Figure 4B, Supplementary Figure S4C). This trend is likely due to species composition differences across niches rather than niche-specific variations within species (e.g., urogenital *E. coli* vs. gut *E. coli*) (Supplementary Figure S4D and E).

**Figure 4.**
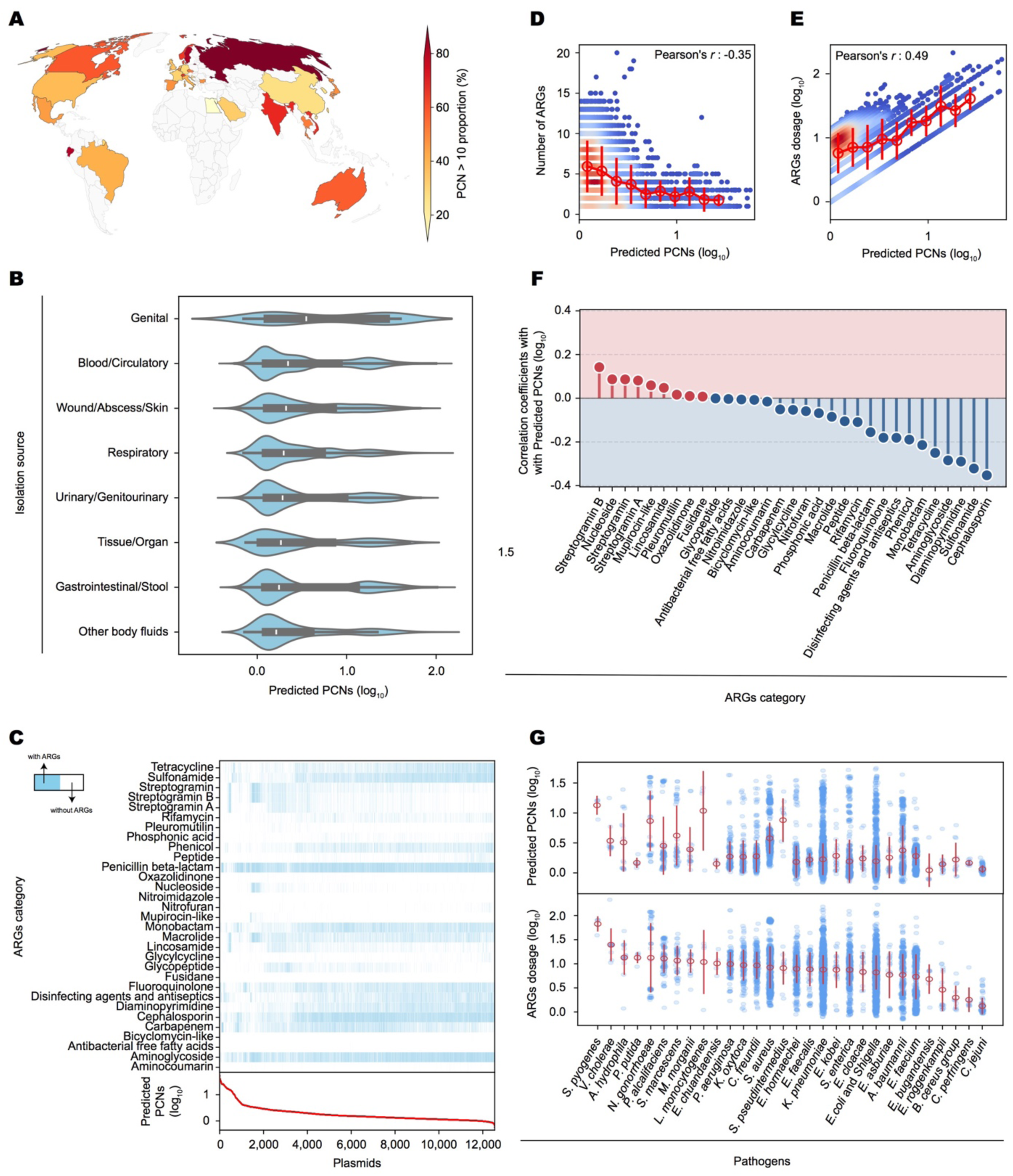
Distribution patterns of predicted PCNs and ARG dosage in clinical plasmids. (A) Geological map showing the proportion of plasmids with PCN > 10 in different countries. (B) Distribution of predicted PCNs across various human isolation sources. The box plot inside each violin represents the median and interquartile range (IQR) of predicted PCNs. The median line for each group is highlighted within the box plot. Groups are sorted by the median predicted PCN in descending order. (C) Distribution of different ARGs in clinical plasmids. The rows represent different ARG categories, while the columns represent individual plasmids. The red curve at the bottom indicates the predicted PCNs across plasmids. (D) Correlation between predicted PCNs and the number of ARGs per plasmid. Color shading represents point density. The PCN range was divided into equal-width bins. Bar plots represent the mean ± standard deviations of ARG numbers per plasmid within each bin. (E) Correlation between predicted PCNs and ARG dosage. Color shading represents point density. The bar plots represent the binned average ± standard deviations. (F) The point-biserial correlation coefficient between predicted PCN and presence/absence of the ARGs per plasmid. Red and blue lines represent positive and negative correlations, respectively. ARG categories are sorted by correlation coefficient. (G) Distribution of predicted PCNs and plasmid ARG dosages across various pathogens, with error bars representing the standard deviation within each group.

To examine the association between PCN and ARG carriage, we annotated ARGs in all pathogenic plasmids, identifying 12,546 plasmids harboring ARGs from 31 drug classes^43^ (Figure 4C, Supplementary Figure S5A; Methods). The prevalence of different ARG categories varied across isolation sources, with penicillin, beta-lactam, and aminoglycoside resistance genes being most common in most clinical sample types (Supplementary Figure S5B). We then calculated ARG richness and dosage for each plasmid, with dosage defined as richness multiplied by PCN. High-copy plasmids showed lower ARG richness (Pearson’s *r* = -0.35, Figure 4D), consistent with their typically smaller sizes. Conversely, ARG dosage correlated positively with PCN (Pearson’s *r* = 0.49) (Figure 4E), suggesting that while high-copy plasmids carry fewer distinct resistance genes per plasmid, their elevated copy number compensates for size constraints, amplifying overall ARG abundance. Analyses across body sites confirmed these trends (Supplementary Figure S5C and D).

While overall ARG richness exhibited a negative correlation with PCN, ARGs from different drug classes showed various correlations. Most ARG types, especially those related to cephalosporin, sulfonamide, diaminopyrimidine and aminoglycoside, displayed negative correlations (Figure 4F). In contrast, streptogramin-type ARGs showed weak positive correlations, indicating that these genes tend to be enriched in small high-copy-number plasmids.

Among different pathogens, *Streptococcus pyogenes*, which causes impetigo, pharyngitis, cellulitis and acute Rheumatic fever, stands out with the highest mean plasmid ARG dosage^44^ (Figure 4G). *Vibrio cholerae*, the causative agent of cholera, and *Aeromonas hydrophila*, which causes gastroenteritis and wound infections, also rank highly^45,46^. These pathogens may pose greater risks in the evolution and dissemination of ARGs. In contrast, *Enterococcus* and *Campylobacter* plasmids display low PCN and ARG dosage, potentially due to chromosomal integration of resistance genes or ecological niches with reduced selection for plasmid-mediated resistance. These findings offer critical insights for risk evaluation and ARG surveillance in prokaryotic pathogens.

### PCN landscape across ecosystems

To investigate PCN distribution across different microbial ecosystems, we further applied our machine learning framework to the IMG/PR dataset, which contains the most extensive collection of plasmid sequences and associated metadata from diverse environments^14^. As many plasmids in the database were derived from metagenomes without validated host organisms, we used the chromosome-independent model relying solely on plasmid-derived features (domains, plasmid *k*-mers, and plasmid length) (Supplementary Figure S6A). For robust analysis, we restricted our predictions to the 136,195 fully sequenced plasmids labeled as ‘putatively complete’.

Predicted PCNs followed a power-law relationship with plasmid size, with a scaling coefficient of –0.90, closely matching that from the training dataset (Supplementary Figure S6B). Our analysis revealed that while engineered ecosystems (wastewater treatment plants, bioreactors) contributed the largest fraction of plasmids (60% of the total), animal-associated environments showed the highest proportion (≈ 80%) of high-copy-number plasmids (PCN > 10). In contrast, plasmids from plant-associated and terrestrial niches were predominantly low-copy (PCN < 10), with mean copy numbers of ∼10 and 13, respectively (Figure 5A). Taxonomic stratification showed PCN variations across prokaryotic classes. Among engineered plasmids (Figure 5B), *Actinomycetia* had the highest median PCN (≈20 copies), followed by *Coriobacteriia* (≈ 18 copies) and *Gammaproteobacteria* (≈ 15 copies). At the low end, *Bacilli* (≈ 10 copies) and *Fusobacteriia* (≈ 8 copies) had the lowest median copy numbers. This pattern also held for plasmids in human gut microbiome (Figure 5C).

**Figure 5.**
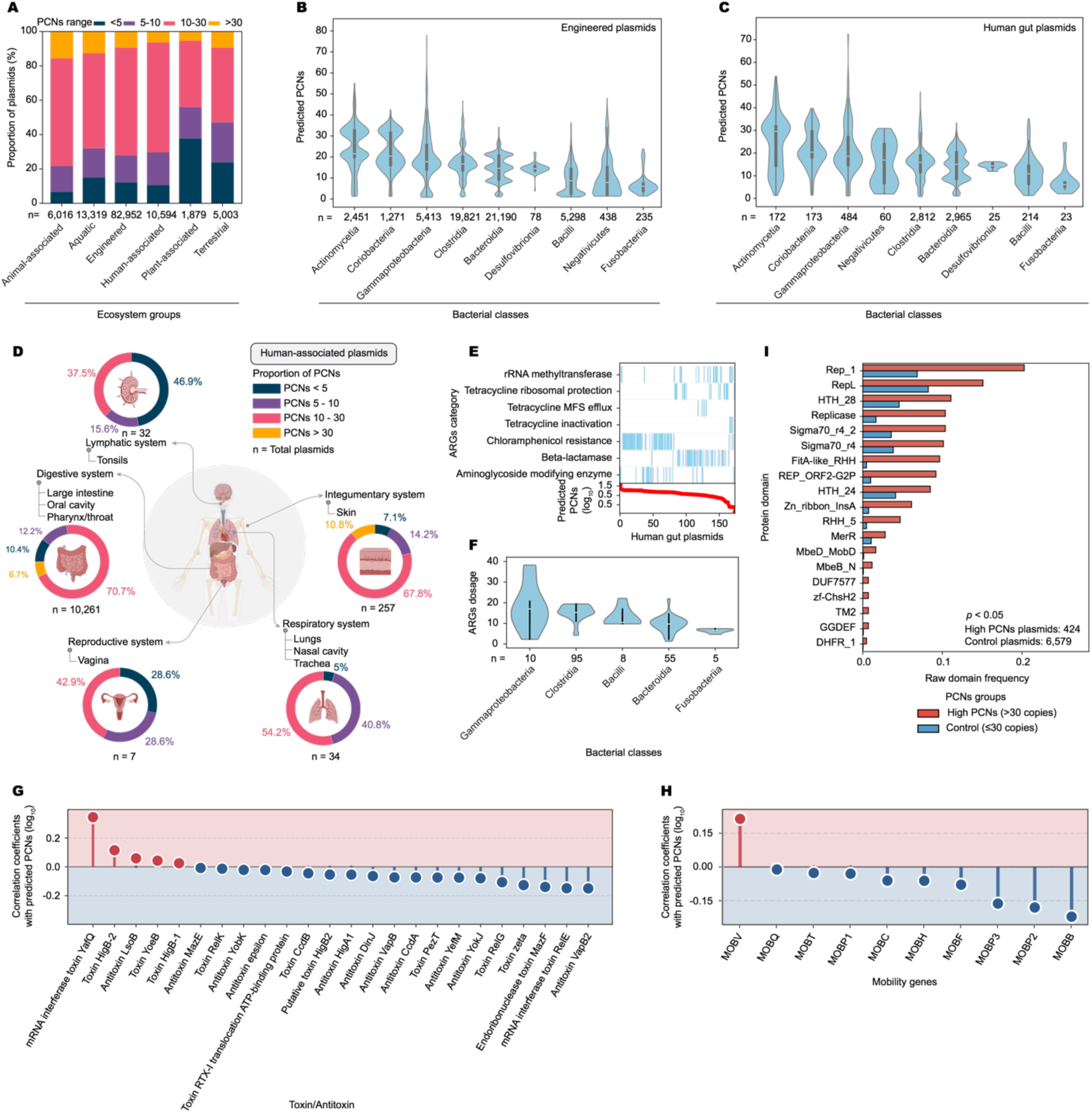
Predicted PCN distribution patterns across different ecosystems. (A) Proportion of plasmids within different PCN ranges in various ecosystem groups. (B) Taxonomic distribution of predicted PCNs in engineered environment, with *n* standing for sample size. (C) Taxonomic distribution of predicted PCNs in human gut microbiome. (D) Pie charts showing the proportion of plasmids within different PCN ranges in different human body systems. (E) Distribution of ARGs across plasmids in human gut microbiome. Rows represent different ARG categories, and columns represent individual plasmids. The red curve at the bottom shows the overall distribution of predicted PCNs across plasmids. (F) Distribution of ARG dosage in human gut plasmids across bacterial classes. Boxplots indicate the median and interquartile range, with *n* representing the sample size. (G, H) Stem plots illustrating the correlation between toxin/antitoxin genes (G) and mobility genes (H) with predicted PCN. Point-Biserial correlation was used for correlation calculation. (I) Frequency of protein domains in high-copy plasmids (PCN > 30, shown in red) compared to other plasmids (PCN ≤ 30, shown in blue).

In human-associated environments, PCNs varied significantly across anatomical sites. The digestive system (the largest plasmid source in human) and integumentary systems emerged as the epicenter of high-copy plasmids, with 77.4% and 78.6% exceeding PCN >10, respectively (Figure 5D). Conversely, the lymphatic system was predominantly low-copy, with 62.5% of plasmids below PCN 10. These patterns likely reflect the niche-specific effects across body sites.

Human gut microbiome is a major reservoir of ARGs^47^. Multiple types of ARGs were detected in the human gut plasmidome (Figure 5E). Chloramphenicol resistance loci are primarily found on mid-to-high copy plasmids. By contrast, β-lactamases are mainly present on low-copy-number plasmids. When stratified by bacterial class, *Gammaproteobacteria* carry the highest ARG dosage, followed by *Clostridia* and *Bacilli* (Figure 5F).

Further analysis identified the associations between PCN and other plasmid traits such as toxicity and mobility (Figure 5G and H). Toxins are the primary pathogenicity factors produced by many bacteria^48^. Many toxins are paired with antitoxins that counteract their toxic effects. Our analysis suggested that in human gut microbiome, many toxin-antitoxin related genes, especially mRNA interferase toxin *yafQ*, were associated with PCN^49^ (Figure 5G, Supplementary Figure S6C). Analysis of conjugation systems also revealed complex relationships with PCN. While most mobility genes showed neutral or negative associations with predicted PCN, the MOBV family demonstrated a strong positive correlation (Pearson’s *r* = 0.20) (Figure 5H, Supplementary Figure S6D). These findings indicate different conjugation systems have evolved for plasmids with different copy numbers.

Many plasmids in human gut microbiome have very high copy numbers (PCN > 30). To understand the mechanisms underlying their high copy number, we compared the protein domains encoded by these plasmids with those encoded by other plasmids. Our analysis revealed two key functional shifts. The replication machinery showed notable enrichment, featuring Replicase domains (6.2-fold increase) along with Rep_1 (2.9-fold; GO:0003677 DNA binding, GO:0006260 DNA replication) and RepL (1.8-fold; GO:0006260 DNA replication, GO:0006276 plasmid maintenance). Transcriptional regulation domains were also prominent, with Sigma70_r4_2 (2.89-fold; GO:0003677 DNA binding, GO:0003700 transcription factor activity, GO:0016987 sigma factor activity) and Sigma70_r4 (2.63-fold; GO:0003700 transcription factor activity, GO:0006352 transcription initiation, GO:0006355 transcriptional regulation)^36^ showing strong enrichment (Figure 5I). This unique functional profile suggests high-copy plasmids might have evolved gut-specific strategies to tightly regulate replication initiation in response to fluctuating nutrient availability and rapidly modulate gene expression to adapt to host dietary changes or immune pressures. These mechanisms not only enhance plasmid persistence in the gut ecosystem but may also facilitate the spread of adaptive traits, including antibiotic resistance genes, among gut microbiota.

## Discussion

Our study integrates theory and machine learning to develop a generalizable framework for predicting PCN. The theoretical model, while intentionally simple, captures how PCN is shaped by multi-level selections and offers a plausible explanation for the empirically observed power-law relationship between PCN and plasmid size. However, this model necessarily overlooks many complexities, such as the functional diversity of plasmid-encoded proteins. Incorporating such factors into a cohesive mechanistic framework is challenging, as the combinatorial parameter space quickly becomes intractable. To address this limitation, we developed a data-driven approach that explicitly incorporates heterogeneous plasmid features. This framework significantly improved prediction accuracy, with plasmid-encoded domains emerging as key PCN predictors. Our machine learning framework complements mechanistic modeling by bridging biological complexity with computational efficiency, thus enabling rapid and large-scale copy number predictions of plasmids sampled from microbiomes across the world. In addition, our framework provides a null baseline for the expected plasmid copy numbers. Measured plasmid copy numbers that significantly exceed this null baseline would evidence strong selection for exceptional plasmid copy numbers in particular contexts, such as very recent exposure to strong antibiotic dosages or other stresses^9,50^.

The performance of our machine learning framework is inherently constrained by the noise in the training dataset^3,19^. For instance, PCN estimations rely on coverage-based methods, which intrinsically introduce more errors for small plasmids. Additionally, the dataset is skewed, with *Enterobacteriaceae* (43%) and human-associated environments (37.7%) overrepresented, while archaea (0.23%) and engineered ecosystems (0.2%) are underrepresented, limiting extrapolation. These biases could be addressed in future works as the validated PCN dataset expands.

Environmental factors can influence PCN, too. For instance, growth conditions (such as temperature, pH and nutrients) can alter the copy number of plasmids^51,52^. PCN can also vary as bacteria transition from logarithmic to stationary growth phase^53^. Additionally, PCN heterogeneity can arise even in clonal populations^11^. The same plasmid may exhibit slightly different copy numbers in different hosts, further complicating PCN prediction^54^. In this work, we relied on sequence features for prediction, while disregarding the confounding factors listed above due to their absence in published sequencing data and a lack of standardized growth conditions across thousands of samples worldwide. This limitation may contribute to the discrepancies between predicted and actual PCNs. Nevertheless, the relatively high accuracy of our machine learning predictions indicates that our approach identifies the most critical features governing PCN control, highlighting the fundamental role of plasmid sequence in determining copy number.

While plasmids are well-known as main carriers of ARGs, the relative contribution of small, high-copy plasmids versus large, low-copy plasmids to ARG dissemination remains unclear. Our method enables the estimation of ARG dosage per plasmid, facilitating risk assessment for plasmid-mediated ARG dissemination. Our results indicate that although high-copy plasmids harbor fewer ARGs due to the size-copy number tradeoff, their elevated copy number may compensate for size limitations, potentially leading to higher overall ARG abundance. This suggests that small, high-copy plasmids may amplify overall ARG dosage, while large low-copy plasmids act as key hubs for ARG dissemination^50,55^. These findings provide key insights for the surveillance, control, and reversal of plasmid mediated ARG transfer.

## Materials and Methods

### Data collection and processing

We analyzed a comprehensive dataset comprising 11,338 fully assembled plasmid sequences from 4,317 prokaryotic genomes with computationally determined copy number estimates from Maddamsetti et al.^3,19^. The PCNs were quantified using the Probabilistic Iterative Read Assignment (PIRA) method, which combines pseudoalignment with iterative refinement for accurate PCN estimation across large sequencing datasets. Following their pipeline, we retrieved complete plasmid assemblies and associated metadata using NCBI Datasets command-line tools, retaining only those entries labeled as “plasmid”. Most plasmids are from natural environments, while some are explicitly labeled as ‘engineered’. These plasmids were constructed or modified in laboratories. Taxonomic information of plasmid hosts are also provided in the associated metadata.

### Feature extraction

For each plasmid in the dataset, we first calculated the plasmid length and, when available, the chromosomal length of the host genome. To characterize sequence composition, we also calculated plasmids *k*-mer frequencies for *k* = 1, 2 and 3. In order to derive the functional features, protein-coding sequences of the plasmids were identified using Prodigal (v2.6.3)^33^. Domain annotation of these protein coding sequences was performed against the Pfam database (v37.1) using HMMER’s HMMScan (v3.4)^56^, applying stringent E-value thresholds (sequence-level: ≤ 0.001; domain-level: ≤ 0.001), to ensure high-confidence domain annotations. This process identified 11,533 unique Pfam domains. Plasmids without detectable domains were excluded from further analysis (retaining 11,051 plasmids), and each domain was represented as a binary feature indicating its presence (1) or absence (0) for the downstream analysis.

To identify domains significantly associated with PCN, we calculated point-biserial correlation coefficients between each binary domain feature and PCN. We adjusted for multiple testing using the Benjamini-Hochberg false discovery rate correction, retaining only domains with q-values < 0.05. This statistical filtering reduced the genomic feature set to 1,288 domains (approximately 9% of the total), which we subsequently characterized functionally using Pfam-to-GO mappings^36^ (version: 2025/04/29 15:50:00) to investigate their potential biological roles in plasmid maintenance and replication.

### Development of machine learning framework

To establish an optimal framework for PCN prediction, we first evaluated the relative contributions of different feature combinations through a tiered approach (Figure 2A and B). We trained parallel models across all possible combinations of four feature categories: (1) the 1,288 statistically significant Pfam domains, (2) absolute *k*-mer frequencies, (3) plasmid length, and (4) host chromosomal length where available. Recognizing the right-skewed distribution of raw PCN values, we applied a log1p transformation (natural logarithm of 1 + PCN) to PCNs. All predicted values were subsequently back transformed to calculate PCNs.

Each combination was independently replicated three times using distinct random seeds to ensure robustness against stochastic training variability. Models’ performance was evaluated using cross-validated R², MSE and Spearman’s correlation values on held-out test (4:1 split) sets, enabling direct comparison between simplified single-feature models and comprehensive multi-feature integrations.

We pre-evaluated various regression algorithms using default parameters: including scikit-learn v1.5.2 simple linear regression^57^, random forest regressor^57^, and XGBoost (v2.1.1)^58^. Consistent preliminary results demonstrated random forest’s superior performance across all feature combinations, leading to its selection as our base algorithm. Subsequent hyperparameter tuning employed a two-phase strategy: an initial broad random search cross validation across wide parameter ranges (documented in code repository), followed by focused grid search cross validation around optimal candidates identified during the exploratory phase.

Practical considerations guided the development of two specialized predictors: (1) a full context model incorporating all four feature types (domains, *k*-mers, plasmid length, chromosomal length) for scenarios with available host genomic data; (2) a plasmid-centric model utilizing only plasmid-derived features (domains, *k*-mers, plasmid length) for broader applicability in metagenomic contexts. In both models, the target variable (PCN) was log1p transformed. Both architectures demonstrated high accuracy (R² ≈ 0.72 and 0.71 respectively), with complete implementation details and datasets publicly available at our github repository.

### Network analysis

To systematically investigate relationships between plasmid-encoded protein domains and copy number variation, we implemented a multi-stage computational pipeline. First, we quantified pairwise similarity between all plasmids based on their binary-encoded domain profiles, where each plasmid was represented as a vector indicating the presence (1) or absence (0) of specific protein domains. We computed cosine similarity scores of all plasmid pairs using scikit-learn’s implementation, which measures the cosine of the angle between vectors to determine their similarity in domain composition while inherently normalizing for differences in plasmid size and domain richness. This produced a symmetric similarity matrix where values ranged from 0 (completely dissimilar) to 1 (identical domain composition).

For network analysis, we established a conservative similarity threshold of ≥0.5 to focus on biologically meaningful relationships, constructing an undirected weighted network where nodes represent plasmids and edges connect plasmids meeting this similarity criterion. The network was imported into Cytoscape (v3.10.3) for visualization^59^, with nodes colored by PCN values using a continuous gradient scale to enable visual assessment of PCN distribution patterns.

To identify functionally related plasmid clusters, we applied the Louvain community detection algorithm to this network^40^. This method iteratively optimizes modularity, grouping plasmids into communities that maximize intra-cluster connections while minimizing inter-cluster edges, resulting in the identification of 112 communities. For downstream analysis, we focused on the five largest communities, comparing their genetic composition and PCN distributions to investigate potential relationships between domain content and copy number variation. Throughout this process, we maintained a clear separation between the quantitative network analysis (performed programmatically) and qualitative visualization (conducted in Cytoscape), ensuring methodological rigor while enabling biological interpretation of the complex network patterns.

### Applications to clinical plasmids

To assess the copy number distribution of clinically relevant plasmids, we obtained genomic data from NCBI’s Pathogen Detection resource on May 02, 2025. We applied stringent filters to select only “complete” genomes from “clinical” isolates, yielding 15,855 genomic accessions. Using NCBI command-line tools, we downloaded these genomes and extracted plasmid sequences by filtering for records explicitly annotated as ‘plasmid’. This process identified 30,254 plasmids from 10,664 genome assemblies, with the remaining genomes containing no detectable plasmids.

To extract the features required for our machine learning framework, we processed these clinical plasmids through an identical bioinformatics pipeline. Protein-coding sequences were predicted using Prodigal, followed by domain annotation with HMMScan against the Pfam database, same versions as of model training. Of the initial 30,254 plasmids, 30,246 contained coding sequences, and we successfully identified at lease one domain in 29,490 plasmids. The convergence between clinical plasmids (8,960 unique domains) and our training feature set, with 1,281 of 1,288 statistically significant domains (99.5%) reappearing in clinical isolates, demonstrates that our feature selection captures fundamental, evolutionarily conserved plasmid elements. For PCN prediction, we constructed a feature matrix comprising plasmid lengths, host chromosomal lengths, absolute *k*-mer frequencies (*k* = 1, 2 and 3), and 1, 288 domains. The full context model generated predictions for log1p-transformed plasmid copy numbers (PCN), which we subsequently back-transformed to PCN values.

To investigate the links between copy number and ARG carriage, we performed comprehensive ARG annotation using the Comprehensive Antibiotic Resistance Database (CARD v4.0.0) and Resistance Gene Identifier (RGI v6.0.3)^43^. This analysis identified ARGs in 12,546 plasmids spanning 31 drug classes, enabling subsequent dose-effect analyses between predicted PCN and resistance gene carriage.

To analyze PCN distribution across isolation sources, we categorized the 18,808 plasmids with clear isolation metadata into broad anatomical groups based on source keywords. The largest groups included: Blood/Circulatory (5,380 plasmids), Gastrointestinal/Stool (4,795), and Urinary/Genitourinary (3,799). A large proportion of remaining plasmids (10,682) either lacked clear source information or originated from non-human/animal hosts. Grouping criteria were systematically defined (e.g., ’Gastrointestinal/Stool’ included terms like ’feces’, ’rectal swab’, and ’intestinal microflora’; ’Urinary/Genitourinary’ encompassed ’urine’, ’UTI’, and ’urethral swab’). This categorization enabled streamlined analysis of PCN patterns across major human anatomical sites while reducing metadata complexity.

To characterize PCN distribution around globe, we analyzed 28,074 plasmids with country-level isolation metadata (spanning 110 countries). Applying a minimum threshold of 100 plasmids per country to ensure statistical reliability, 31 countries qualified for final analysis. High-PCN prevalence was calculated per country by dividing the count of plasmids with PCN>10 by the total plasmids isolated from that region. Geospatial mapping was implemented using Natural Earth’s administrative boundaries (1:110m scale, v5.1.1; downloaded from https://www.naturalearthdata.com) projected in Robinson projection via GeoPandas.

### Application to IMG/PR dataset

We implemented a comprehensive analytical pipeline to apply our model to plasmids in the IMG/PR database. Beginning with data acquisition, we programmatically retrieved all available plasmid sequences and their associated metadata through the JGI Data Portal API (https://genome.jgi.doe.gov/portal/IMG_PR). The collected metadata included essential information such as plasmid topology, ecosystem, host taxonomy, plasmid length, and functional annotations including mobility genes and antibiotic resistance determinants. To ensure data quality, we established stringent filtering criteria. We excluded sequences containing more than 5% ambiguous bases and removed all plasmids shorter than 1 kb. Additionally, we only selected those plasmids labeled as “putatively complete”. This quality control process yielded a final dataset of 154,680 high-quality plasmid sequences from an initial collection of 699,978 entries, providing a robust foundation for downstream analysis.

The feature extraction process precisely followed our established training methodology while incorporating necessary adaptations for metagenomic data. We implemented a custom Python script for *k*-mer frequency calculation. For protein-coding sequence prediction, we utilized Prodigal v2.6.3 in metagenomic mode (-p meta), specifically optimized for the fragmented and heterogeneous sequences characteristic of environmental samples. The predicted coding sequences were analyzed using the same pipeline as used in the model, identifying domains in 136,195 plasmids that met quality standards.We assessed domain conservation across plasmid populations in the IMG/PR dataset. Human gut plasmids had 70.3% of reference domains, engineered plasmids had 98.8%, and environmental plasmids showed 99.8%. These high overlap rates validate our feature selection and support the model’s broad applicability. The final implementation of our plasmid-centric model for PCN prediction across these diverse ecosystems maintained consistency with our training approach, using log1p-transformed values that were subsequently back-transformed.

Plasmids were categorized into seven ecosystem groups based on metadata annotations: Terrestrial, Engineered, Plant-associated, Aquatic, Animal-associated, Human-associated, and Microbial-associated (excluded in the following analysis due to ambiguous annotations). We employed a hierarchical classification approach, prioritizing specificity—for instance, by segregating human-associated plasmids. Subcategories (e.g., “Engineered,” encompassing bioreactors and wastewater systems) were established to enhance clarity. Plasmids that did not align with predefined categories were classified as “Uncategorized”.

To identify toxin/antitoxin related genes in the plasmids from the human gut, we performed Prokka annotation, followed by searching for the term “toxin/antitoxin” in the product descriptions. Only plasmids containing toxins or antitoxins were retained for further analysis.

We compared protein domain frequencies between high-PCN plasmids (>30 copies) and control plasmids (≤30 copies) using one-tailed Mann-Whitney U tests to identify domains enriched in high-PCN plasmids. To account for multiple comparisons, we applied Benjamini-Hochberg false discovery rate (FDR) correction, with statistical significance defined as FDR-adjusted p< 0.05. Enriched domains were further filtered to include only those showing both statistical significance (FDR p < 0.05) and biological relevance (fold-enrichment > 1 in high-PCN plasmids). For visualization, we focused on the top 20 most enriched domains (ranked by raw domains frequency) to highlight the strongest associations with high copy number maintenance.

## Data Availability

All the data associated with this work are available at the Github repository (https://github.com/Iqra123isynbio/Plasmid_copy_number_Prediction).

## Code Availability

All codes are available at the Github repository (https://github.com/Iqra123isynbio/Plasmid_copy_number_Prediction).

## Acknowledgement

This study was supported by the National Key R&D Program of China (2024YFA0920200 to TW), the National Natural Science Foundation of China (12401660 and 32470701 to TW), and the Shenzhen Institute of Synthetic Biology Scientific Research Program (HSE499011086 to TW). We are grateful to the Shenzhen Infrastructure for Synthetic Biology for providing instrument support and technical assistance.

