## Supplementary Information for "Integrating theory and machine learning to reveal determinants of plasmid copy number"

**Supplementary Text**

**A simple model explaining the power law relationship between plasmid size and copy number**

To explain the power-law relationship between plasmid size and copy number, we adapted a model describing the intracellular resource competition between the chromosome and plasmids[1]. In this framework, the strengths of host-level and plasmid-level selections are determined by how resources are distributed between the chromosome and plasmids.

Let $C$, $P$ and $\theta$ denote chromosome size, plasmid size and PCN, respectively. Both $C$ and $P$ are assumed to be proportional to the number of genes encoded[1]. We further assume that a larger gene repertoire enhanced the cell's ability to exploit diverse environmental resources, thereby increasing the total intracellular resource availability $R_{total}$. The relationship between genome size and $R_{total}$ is described by a Hill function:

$$R_{total} = \frac{M(C + h\theta P)}{K + C + h\theta P}. \quad (1)$$

Here, $M$ is the maximum attainable resource availability, $h$ is a constant reflecting the relative advantage of plasmid genes in resource competition. $K$ is the value of $C + h\theta P$ at which half of $M$ is reached.

The resources devoted to plasmid functions ($R_P$) are determined by the ratio of plasmid DNA to total DNA:

$$\frac{R_P}{R_{total}} = \frac{h\theta P}{C + h\theta P}. \quad (2)$$

Combining equation (1) and (2) gives:

$$R_P = \frac{h\theta MP}{K + C + h\theta P}. \quad (3)$$

Plasmid-level selection favors greater $\frac{R_P}{R_{total}}$. Therefore, we defined the cost function ($\phi_{plasmid}$) of plasmid-level selection as:

$$\phi_{plasmid} = 1 - \frac{R_P}{R_{total}} = \frac{C}{C + h\theta P}. \quad (4)$$

$\phi_{plasmid}$ can be reduced by increasing $\theta$ or $P$.

Similarly, the resources allocated to host functions ($R_C$, encoded by chromosome) are:

$$R_C = \frac{MC}{K + C + h\theta P}. \quad (5)$$

When the host is plasmid-free, all resources are allocated to the chromosome, giving:

$$R_0 = \frac{MC}{K + C}. \quad (6)$$

Plasmid carriage reduces this allocation to $R_C$, and the different between $R_C$ and $R_0$ represents the metabolic burden of plasmid maintenance. We therefore define the cost function of host-level selection as:

$$\phi_{host} = 1 - \frac{R_C}{R_0}, \quad (7)$$

which simplifies to:

$$\phi_{host} = \frac{h\theta P}{K + C + h\theta P}. \quad (8)$$

Host-level selection thus acts to minimize $\phi_{host}$ by reducing $\theta$ or $P$.

The total cost function is given by the sum of the two components:

$$\phi_{total} = \frac{h\theta P}{K + C + h\theta P} + \frac{C}{C + h\theta P}. \quad (9)$$

Here, PCN ($\theta$) increases $\phi_{host}$ but decreases $\phi_{plasmid}$. Consequently, equation (9) predicts the biphasic change of $\phi_{total}$ with increasing $\theta$: starting from 1, $\phi_{total}$ decreases to a minimum, then rises back toward 1. At low $\theta$, $\phi_{plasmid}$ dominates; at high $\theta$, $\phi_{host}$ dominates. The minimum of $\phi_{total}$ represents the evolutionary optimum of $\theta$.

To obtain this optimum analytically, we calculate the derivative of $\phi_{total}$ with respect to $\theta$:

$$\frac{\partial \phi_{total}}{\partial \theta} = \frac{hKP + hPC}{(K + C + h\theta P)^2} - \frac{hPC}{(C + h\theta P)^2}. \quad (10)$$

The optimal $\theta$, denoted as $\theta^*$, could be derived by solving the equation $\frac{\partial \phi_{total}}{\partial \theta} = 0$:

$$\frac{hKP + hPC}{(K + C + h\theta P)^2} - \frac{hPC}{(C + h\theta P)^2} = 0, \quad (11)$$

which led to

$$\theta^* = \frac{\sqrt{KC + C^2}}{hP}. \quad (12)$$

Equation (12) reveals the power-law relationship between optimal PCN ($\theta^*$) and plasmid size ($P$). Moreover, it predicts that the total plasmid DNA content ($\theta^* \cdot P$) scales positively with chromosome size ($C$):

$$\theta^* \cdot P = \frac{\sqrt{KC + C^2}}{h}. \quad (13)$$

This prediction is consistent with observations from our PCN dataset.

### Supplementary Figures

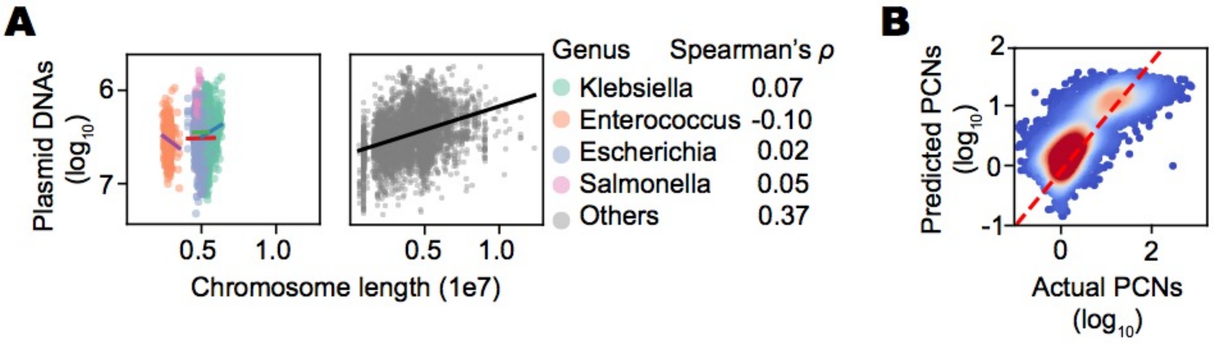


**Figure S1| Plasmid size exhibits limited predictive power for PCN.**

(A) Correlation between plasmid DNA amounts and chromosome size. The four most prevalent bacterial genera (*Escherichia, Klebsiella*, *Salmonella*, and *Enterococcus*) are shown on the left, while other genera are displayed on the right. Linear regressions ($log_{10}$-transformed plasmid DNA vs. chromosome size) are depicted as straight lines.

(B) Correlation ($R^2$ ~0.63) between actual PCNs and predicted PCNs derived from the power-law relationship between plasmid size and PCN. Point density is represented by color shading.

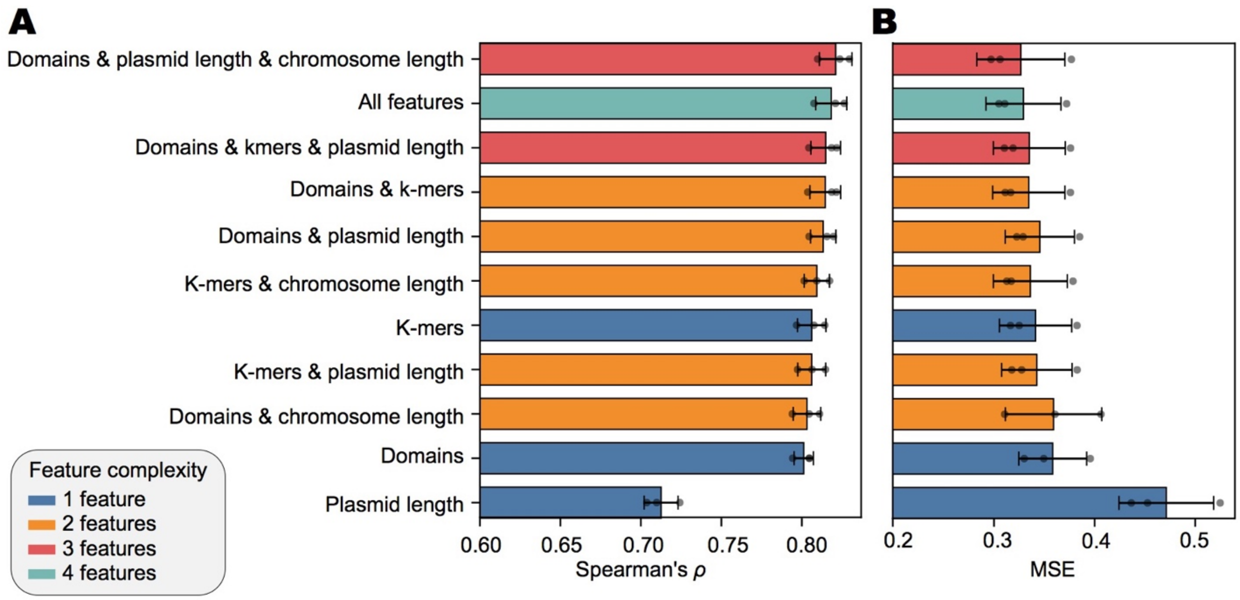


**Figure S2| The predictive performances of the machine learning framework using different sequence-derived features.**

(A) Model performances evaluated by the Spearman's $\rho$ between actual and predicted PCNs. Error bars represent the standard deviations across 3 replicates.

(B) Model performances evaluated by MSE (mean squared error) between actual and predicted PCNs.

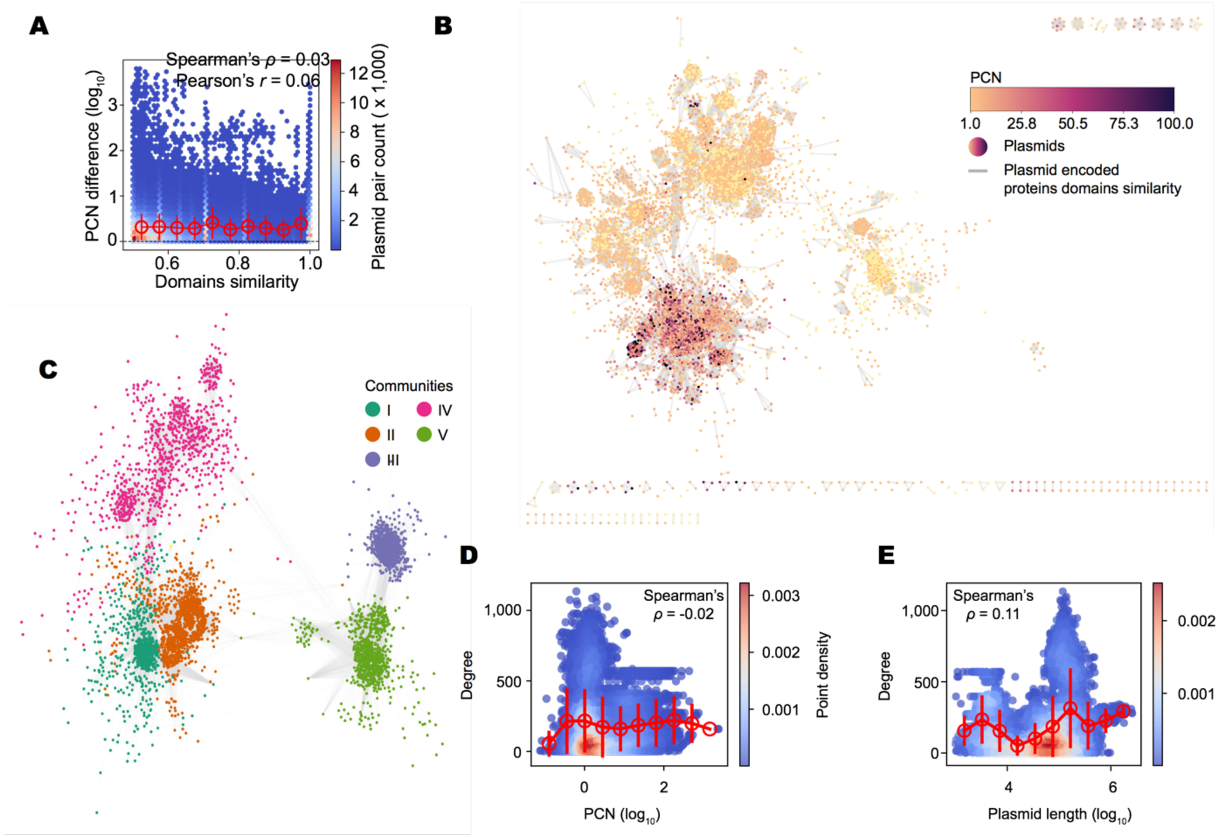


**Figure S3| Plasmid similarity network based on their encoded protein domains.**

(A) Relationship between cosine similarity scores and PCN differences across all plasmid pairs. For each pair, PCN difference is defined as the ratio of the larger PCN to the smaller one. Bar plots represent binned averages ± standard deviations.

(B) Network visualization. Nodes represent individual plasmids, colored by their PCNs. Edges connect plasmids with a cosine similarity of $\geq 0.7$ in their protein domain compositions.

(C) Community structure of the plasmid network. Nodes are colored according to their assigned community membership (I-V).

(D) Association between plasmid degree (connectivity) and PCN. PCN values were grouped into equal-width bins, with bar plots showing the mean degree ± standard deviation within each bin. Degree is defined as the number of plasmids connected to a given plasmid in the network created with the threshold of cosine similarity $\geq 0.5$.

(E) Relationship between plasmid degree and plasmid size.

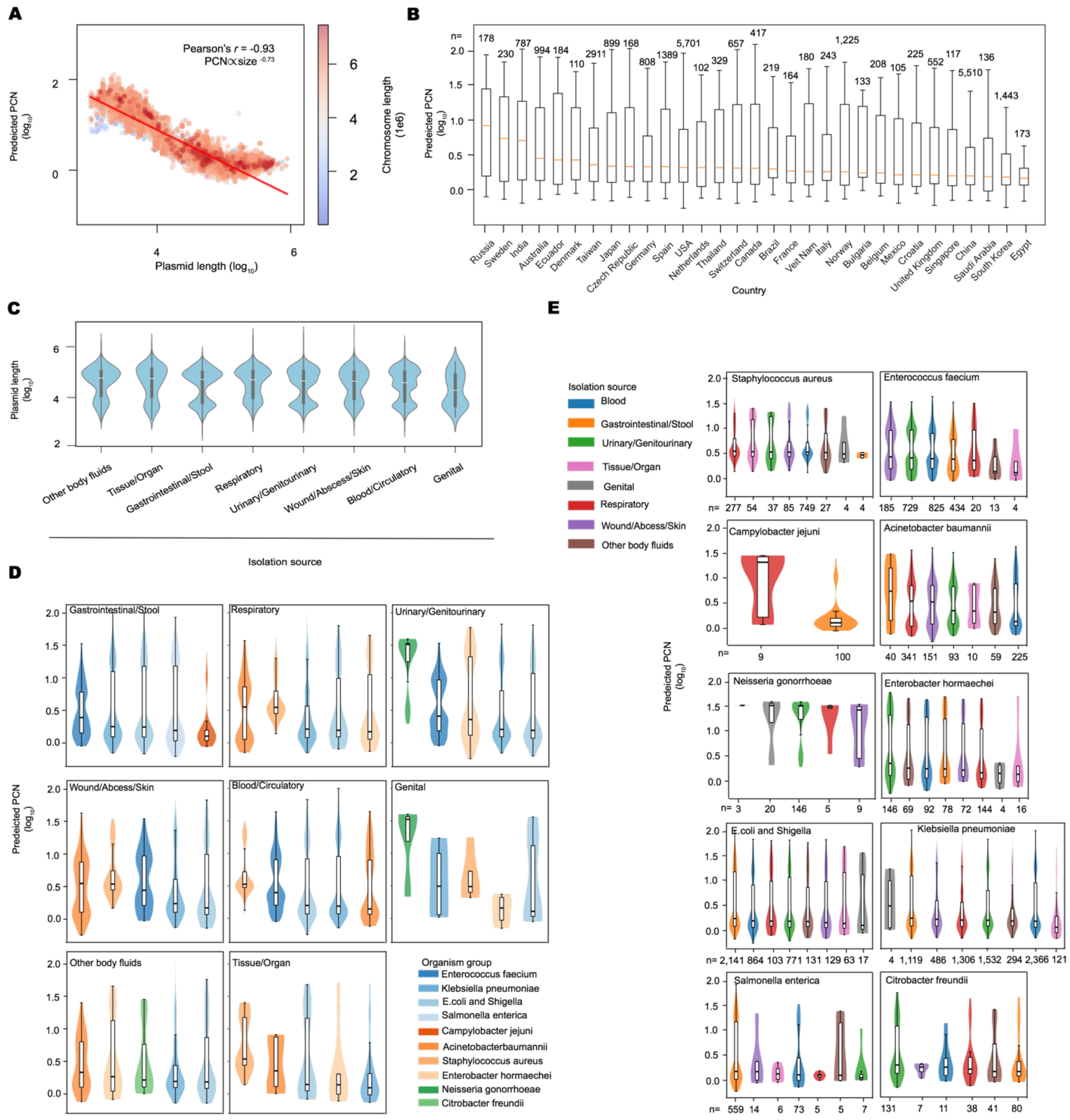

**Figure S4| Distribution patterns of predicted PCNs in clinical plasmids.**

(A) Power-law relationship between predicted PCNs and sizes of clinical plasmids, with a scaling coefficient of -0.73. Point density is indicated by color shading (coolwarm scale).

(B) The distribution of predicted PCNs across different countries. Here, *n* represents the number of clinical plasmids mapped to each country and is shown on the top of each box.

(C) Distribution of plasmid size across various human isolation sources. Groups are ordered by median plasmid length (largest to smallest).

(D) Distribution of predicted PCNs across various organism groups, stratified by human isolation sources. Violin plots show the full distribution, with embedded box plots indicating the median and interquartile range (IQR). Median values are highlighted within each box plot, and groups are arranged in descending order of median PCN.

(E) Distribution of predicted PCNs across various human isolation sources, stratified by organism groups.

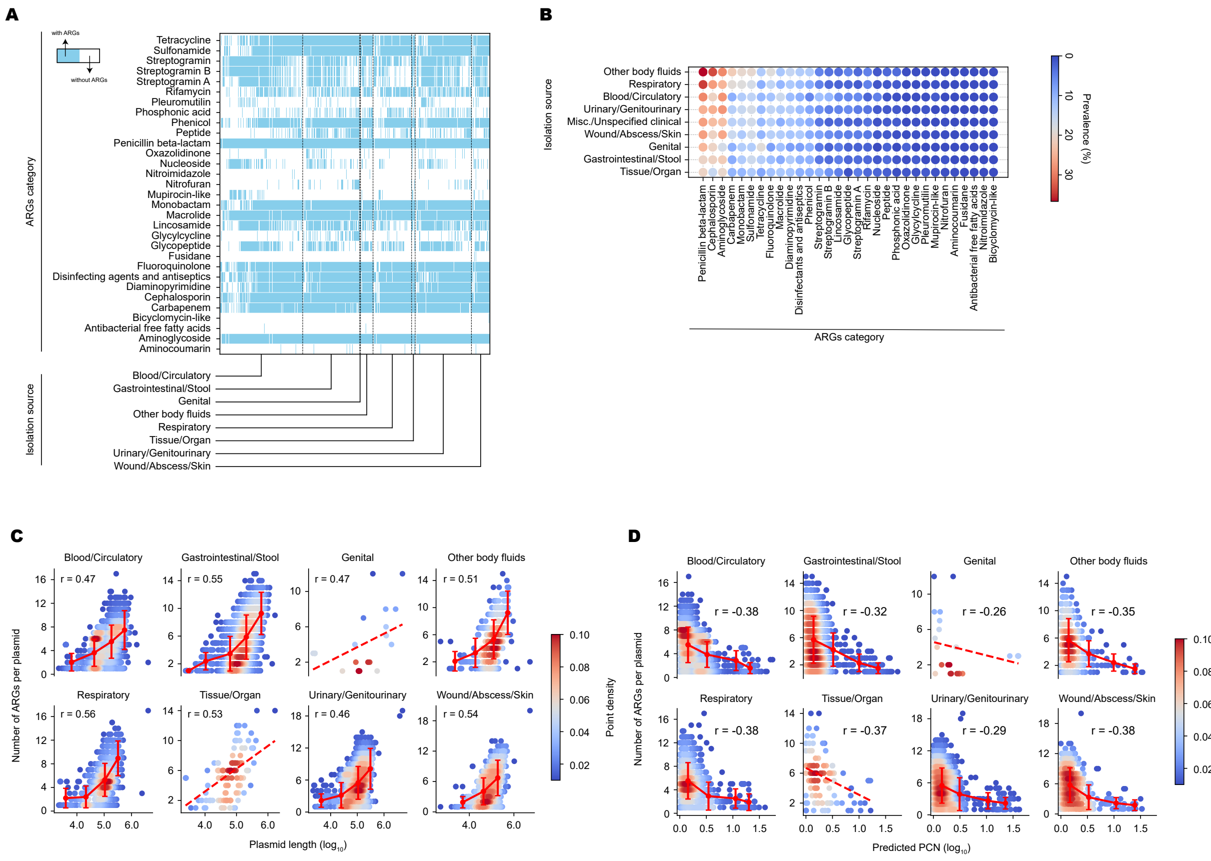

**Figure S5| Distribution patterns of ARGs in clinical plasmids.**

(A) Distributions of 32 categories of ARGs across plasmids in different human body systems.

(B) Prevalence of different ARG types in different human body systems.

(C) Relationship between plasmid length and the number of ARGs per plasmid across body systems. Point density is indicated by color shading, while bar plots show binned averages ± standard deviations. *r* denotes Pearson's correlation coefficient.

(D) Relationship between predicted PCN and the number of ARGs across body systems.

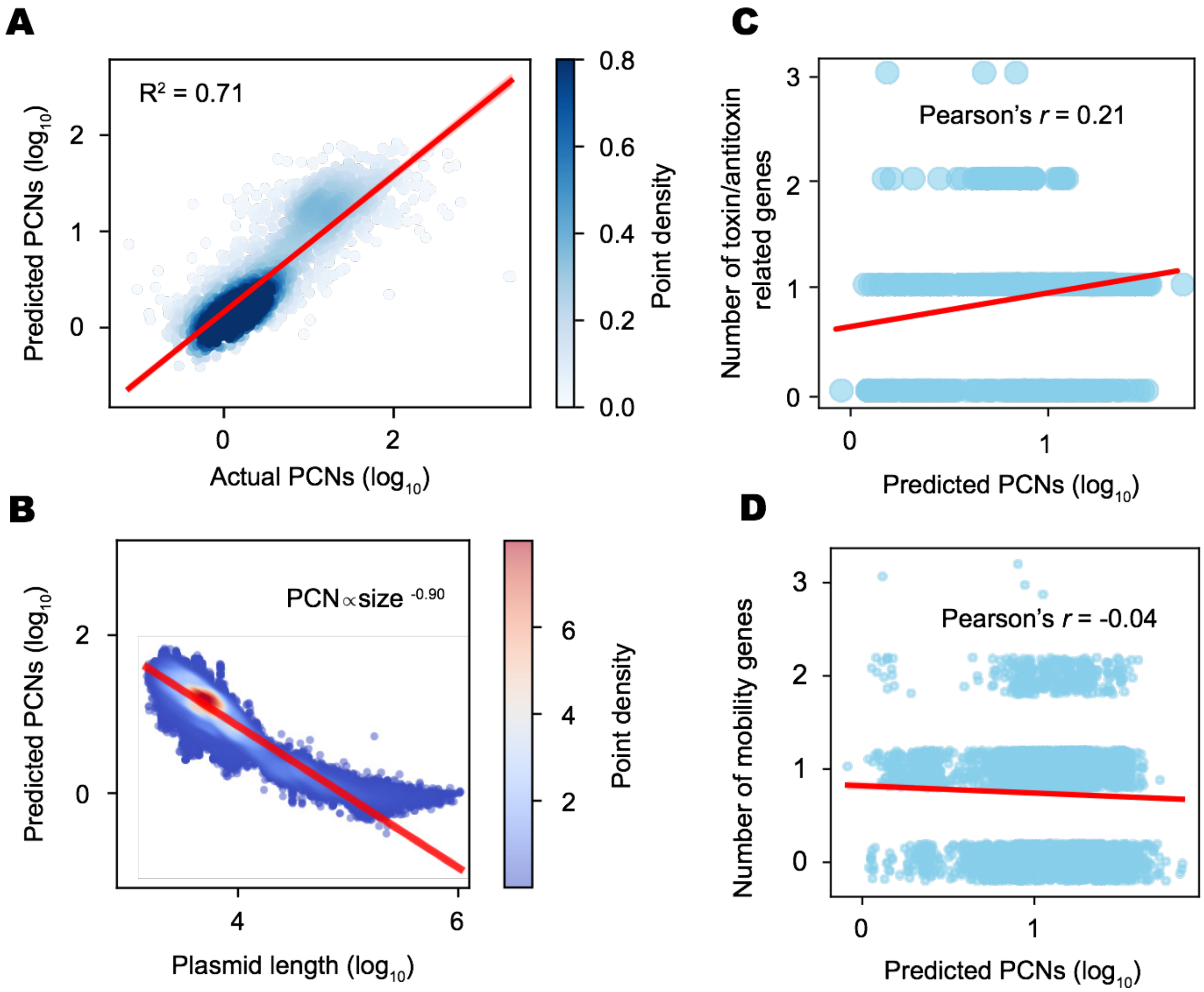


**Figure S6| Distribution pattern of predicted PCNs across ecosystems.**

(A) Correlation between predicted and actual PCNs in IMG/PR dataset. Point density is indicated by color shading (darker indicates higher density).

(B) Power-law relationship between plasmid size and predicted PCN in IMG/PR dataset, with a scaling coefficient of -0.90.

(C) Correlation between predicted PCNs and the number of toxin/antitoxin genes per human gut plasmid. The red line represents the linear regression.

(D) Relationship between predicted PCNs and the number of mobility-related genes per human gut plasmid.
